# Biosynthetic gene clusters in *Pseudomonas viridiflava* have a fitness cost during *Arabidopsis thaliana* infection

**DOI:** 10.64898/2026.02.17.706262

**Authors:** Alejandra Duque-Jaramillo, Efthymia Symeonidi, Manuela Neumann, Haim Ashkenazy, Detlef Weigel, Talia L. Karasov

## Abstract

Specialized or secondary metabolites mediate biotic interactions, including virulence and defense. In plant-pathogenic *Pseudomonas*, certain specialized metabolites can enhance colonization of plant hosts, yet their broader contribution to plant–microbe interactions and the relative importance of different metabolites remain unclear. Specialized metabolites are products of enzymes encoded in biosynthetic gene clusters (BGCs), whose prediction from genome sequences has become routine but whose functional roles are rarely tested experimentally. Here, we characterize the BGC repertoire of 225 *P. viridiflava* isolates from *Arabidopsis thaliana* and assess BGC contributions to fitness *in planta* and disease severity. The BGC landscape of *P. viridiflava* was dominated by non-ribosomal peptide synthetase (NRPS) and NRPS-like BGCs, with one-third of families restricted to a single isolate. Transposon mutagenesis coupled with random barcode transposon sequencing (RB-TnSeq) revealed that the majority of BGCs reduce rather than increase fitness during *A. thaliana* infection, with the magnitude of the fitness cost varying across host genotypes. This cost could be due to exploitation of public goods by cheater mutant strains. In single-isolate plant infections, where public goods are not available, several BGC families were negatively associated with disease severity, which is positively correlated with bacterial growth in this pathosystem, further indicating that BGCs are generally not beneficial *in planta*. Our findings reveal extensive and largely uncharacterized biosynthetic potential in populations of *P. viridiflava* and indicate that candidate metabolites are likely not adaptive for direct interactions with the plant, but perhaps for microbe-microbe interactions either *in planta* or in other ecological niches.

**IMPORTANCE:** Bacteria living on plant leaves produce a vast array of chemical compounds, called secondary or specialized metabolites, that can mediate their interaction with the plant host or other microorganisms. Some of these compounds are known to directly influence how bacteria interact with plants, but it has been unclear whether this is a general rule. We studied a large collection of closely related leaf-dwelling bacteria that varied in their ability to cause disease, focusing on leaf-associated *Pseudomonas viridiflava*—a plant pathogen. We found that very few of the gene clusters responsible for making specialized metabolites improved the ability of the bacteria to colonize *Arabidopsis thaliana*. On the contrary, carrying these gene clusters often reduced bacterial growth and disease severity in plants. Specialized metabolites may instead primarily be important for interacting with other microbes, different host species, or under environmental conditions we did not test. These are questions that remain for future research.

## INTRODUCTION

Bacteria can produce a wide range of specialized metabolites, also known as secondary metabolites, which mediate interactions with other organisms and generally help bacteria to cope with their environment. Specialized metabolites are defined as metabolites that are not essential for basal bacterial growth or division but that instead play a role in nutrient acquisition, quorum sensing, defense and virulence (1). Specialized metabolites also mediate the interaction between bacteria and host organisms they colonize, such as plants (2). Some of these metabolites are well known to affect plant growth and plant defense against pathogenic bacteria. Examples of specialized metabolites that mediate plant-bacterial interactions and contribute to the virulence of plant-pathogenic *Pseudomonas* species include toxins such as coronatine, syringomycin and phaseolotoxin (3), the iron-chelating siderophore pyoverdin (4) and biosurfactants syringafactin and cichofactin, which increase the availability of water-insoluble substrates and promote swarming (5, 6).

Specialized metabolites are produced by proteins encoded in biosynthetic gene clusters (BGCs). A BGC is a consecutive array of at least two genes that encode the biosynthetic pathway for the production of these metabolites and their variants, including biosynthetic, regulatory, and transport proteins (7, 8). BGCs have often defined patterns of different types of genes depending on the chemical class of specialized metabolites they produce; hence, they can be predicted from genome sequences. The gold-standard tool for BGC prediction in bacterial genomes is antiSMASH (<u>anti</u>biotics and <u>s</u>econdary <u>m</u>etabolite <u>a</u>nalysis <u>sh</u>ell) (9, 10), which identifies BGCs based on profile Hidden Markov Models (HMMs) of genes that are specific for each metabolite class. Genome mining for BGCs in small and large bacterial genomes datasets is now common, particularly focusing on a genus or species (11, 12) or on a particular environment, from the human microbiota (13) to oceans (14), soil (15, 16) and many other habitats (17, 18).

*Pseudomonas viridiflava* is a globally-distributed agricultural pest (19–21) and a natural pathogen of *Arabidopsis thaliana* (*22–24*), but its mechanisms of virulence are largely unknown. *Pseudomonas viridiflava* is prevalent in the epi- and endophytic leaf compartments in *A. thaliana* populations in Southwest Germany and across Europe (24, 25). Most isolates are pathogenic on *A. thaliana* under lab conditions, with inter-strain variation in virulence (21, 24, 26, 27). These *P. viridiflava* isolates, referred to as ATUE5 as they were first isolated <u>A</u>round <u>Tu</u>ebingen encode two known virulence factors: the effector AvrE and the pectate lyase gene *pel*, and lack other well-known effectors proteins and toxins (24).

The *Pseudomonas* genus is well-known for its ability to produce a vast array of specialized metabolites (28–30) and it is one of the most biosynthetically-diverse bacterial genera, second only to *Streptomyces* (31). There is great diversity in the specialized metabolite potential of different *Pseudomonas* species (29, 30, 32); yet, variation among isolates of the same species has rarely been studied. Descriptions of the BGC repertoire of plant-associated *Pseudomonas* have been published in recent years, focusing on isolates from the soil and the plant rhizosphere (12, 30, 33, 34). However, most studies did not test the importance of these BGCs in the interaction with the host and/or other microbes (35–37), particularly beyond well-known phytotoxins. Considering that *P. viridiflava* isolates are pathogenic to varying degrees on *A. thaliana* but lack effector proteins and toxins encoded by other plant-pathogenic *Pseudomonas* species, we hypothesized that *P. viridiflava* encodes specialized metabolites that are involved in infection and disease severity on *A. thaliana*.

Here, we describe the largely-uncharacterized specialized metabolite potential of 225 closely-related yet genetically distinct *P. viridiflava* ATUE5 isolates previously obtained from *A. thaliana* populations in Southwestern Germany (24). We not only predict the BGC repertoire from whole-genome sequences but also show that BGC genes reduce fitness during plant infection, with varying magnitude between host genotypes. By integrating the pattern of BGC presence/absence with disease severity data of single-isolate infections of *A. thaliana*, we identify BGCs that negatively correlate with disease severity on plants, further corroborating that they incur fitness costs *in planta*.

## RESULTS

### The biosynthetic potential of *A. thaliana*-associated *P. viridiflava* is broad and its products are largely unknown

We selected representative *P. viridiflava* ATUE5 genomes from a published collection (24) based on gene content. Starting with 1,338 complete genome assemblies (BUSCO single genes > 95%), we reduced our dataset to 163 ATUE5 genomes with distinct gene content according to the presence and absence of groups of orthologous genes. In addition, we included the genomes of 62 ATUE5 isolates for which disease severity data was available, measured as plant size (green pixels) post-infection (27). A total of 225 ATUE5 genomes were included in downstream analyses (Table S1). These genomes had a median completeness of 98.1% (min = 92.1%, max = 99.2%) and 88 contigs (min = 19, max = 5,666) (Table S1). Completeness was not available for four genomes, one of which also lacked the number of contigs in the assembly. Representative genomes had higher completeness (>94%) and smaller number of contigs than genomes with disease severity data.

We characterized the specialized metabolite potential of these 225 *A. thaliana*-associated *P. viridiflava* isolates with antiSMASH 7 (10), predicting their BGCs repertoire. When two or more BGC were predicted in the same genomic region, we counted each BGC independently. In total, 2,925 regions encoding 3,519 BGCs were predicted (Figure 1).

**Figure 1.**
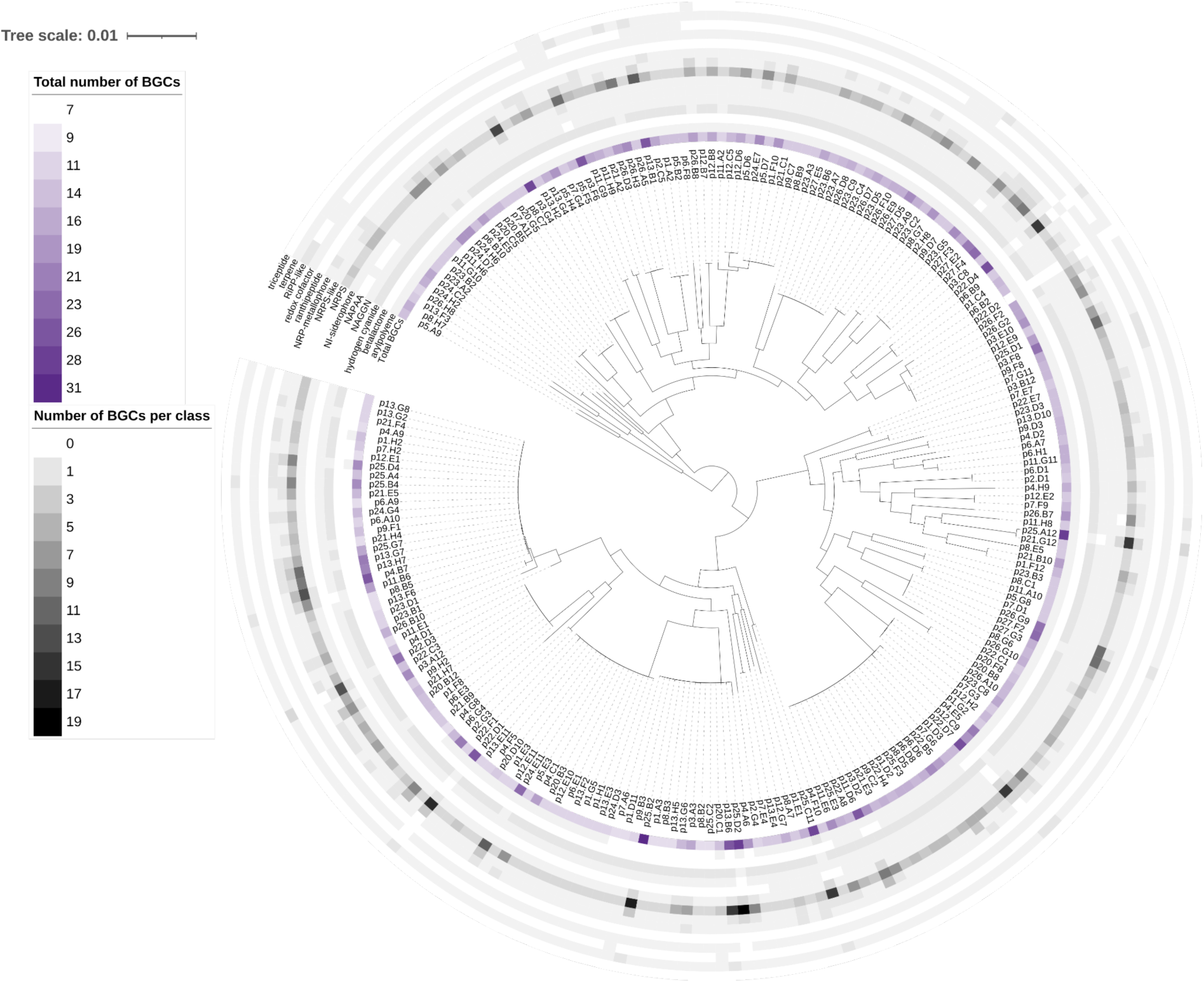
Distribution of biosynthetic gene clusters across 225 genomes of plant-associated *Pseudomonas viridiflava*. Core genome maximum likelihood phylogeny (24) of 225 *P. viridiflava* isolates from Southwest Germany. Innermost ring (purple): total number of predicted BGCs, outer rings (grey): number of BGCs per class, for classes arylpolyene, betalactone, hydrogen cyanide, N-acetylglutaminylglutamine amide (NAGGN), non-alpha poly-amino acids like e-Polylysin (NAPAA), NRP-independent (NI) siderophore, NRPS, NRPS-llike, NRP metallophore, ranthipeptide, redox cofactor, RiPP-like, terpene and triceptide. BGC: biosynthetic gene cluster, NRPS: non-ribosomal peptide synthetase, RiPP: ribosomally synthesized and post-translationally modified peptide.

Genomes had on average 15.6 BGCs (median = 15, min = 7, max = 31; Figure 2A). The percentage of BGCs on a contig’s edge ranged from 0 to 100%, with a mean of 41% and a median of 38%. We visually examined the relation between the total number of BGCs and the percentage of BGCs on a contig’s edge with the assembly statistics (Figure S1). We found that genomes with more contigs and with lower completeness, i.e., of genomes of lower quality, had more predicted BGC in general, and more of the predicted BGCs were located on contig edges (Pearson’s correlation coefficient magnitude between 0.34 and 0.48).

**Figure 2.**
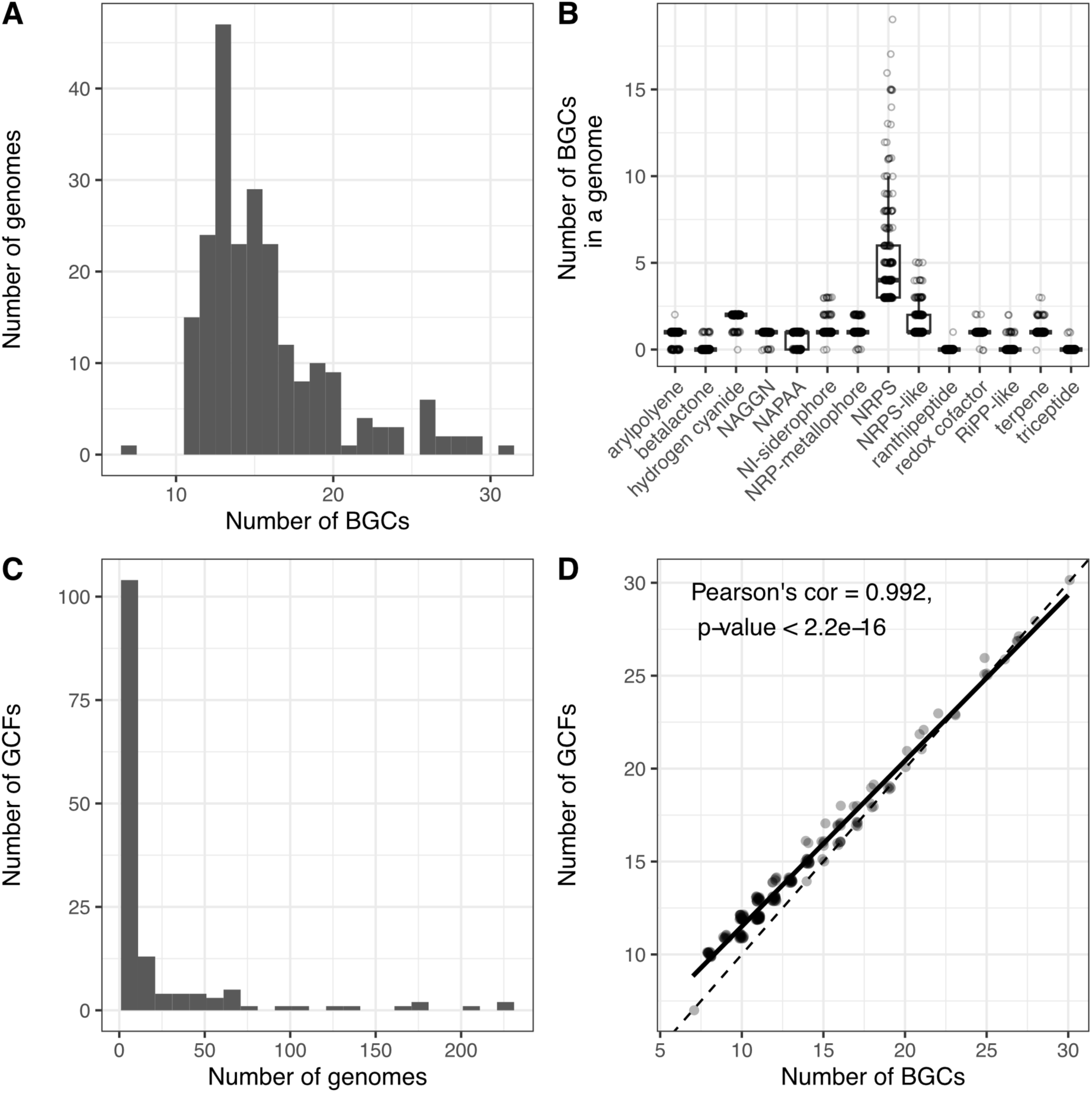
Specialized metabolite potential of *Pseudomonas viridiflava* associated with *Arabidopsis thaliana*. A. Distribution of the number of BGCs predicted per genome. B. Boxplot of the number of BGCs predicted per genome for each detected class. C. Distribution of the number of genomes a given gene cluster family (GCF) is detected in. D. Number of GCFs relative to all BCGs for each genome. Solid line: linear regression of the data, dashed line: unity. BGC: biosynthetic gene cluster (output from antiSMASH), GCF: gene cluster family (output from BiG-SCAPE).

**Figure S1.**
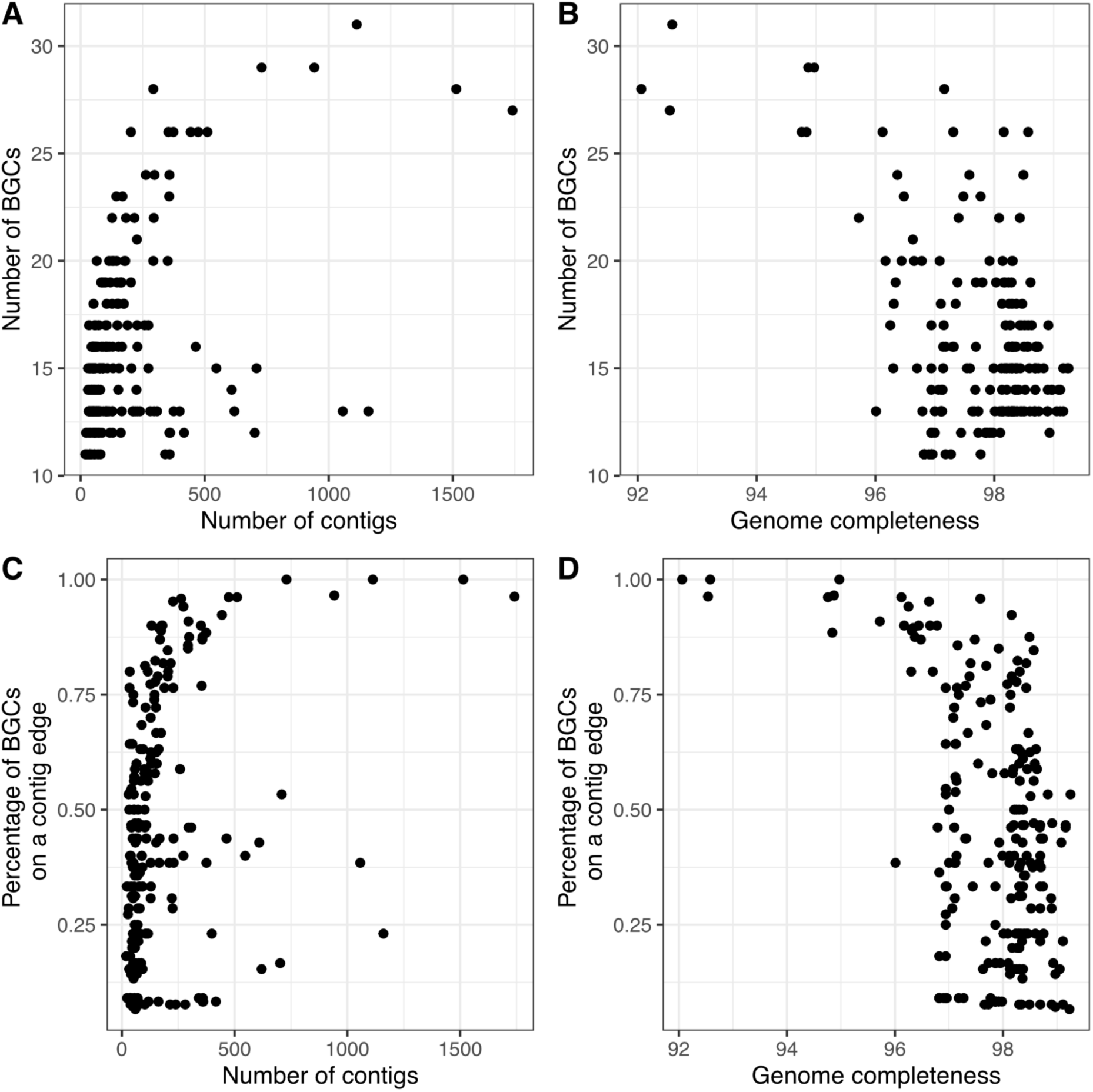
Relation between genome assembly statistics and BGC prediction. The number of predicted BGCs vs the number of contigs in a genome (A) and the genome’s completeness as per CheckM (B). Proportion of predicted BGCs located in a contig edge vs the number of contigs in a genome (C) and the genome’s completeness as per CheckM (D).

The predicted BGCs represented 14 of the 81 (17%) specialized metabolite chemical classes predicted by antiSMASH 7. The most abundant BGC classes were NRPS and NRPS-like, accounting for 43% of all the predicted BGCs (Table 1). These were the only BGC classes predicted in all isolates: every genome had at least three NRPS and one NRPS-like BGC (max = 19 and 5, respectively; Figure 2B, Table 1). BGCs for hydrogen cyanides, NRP-metallophores, NI-siderophores, terpenes, redox cofactors and NAGGNs were found in at least 90% of the genomes; together these classes represented an additional 46% of all the BGCs predicted in 225 ATUE5 genomes. On the other end of the frequency spectrum, BGCs for RiPP-like, betalactones, triceptides and ranthipeptides were predicted in fewer than 10% of the genomes, representing only about 1% of all predicted BGCs.

**Table 1.** Number of predicted biosynthetic gene clusters per class in 225 plant-associated *P. viridiflava* genomes.

| BGC class | n | % of all<br>predicted<br>BGCs | min | max | mean | median |
| --- | --- | --- | --- | --- | --- | --- |
| NRPS | 1,186 | 33.7 | 3 | 19 | 5.3 | 4 |
| Hydrogen cyanide | 426 | 12.1 | 0 | 2 | 1.9 | 2 |
| NRPS-like | 351 | 10.0 | 1 | 5 | 1.6 | 1 |
| NRP Metallophore | 260 | 7.4 | 0 | 2 | 1.2 | 1 |
| NI-Siderophore | 257 | 7.3 | 0 | 3 | 1.1 | 1 |
| Terpene | 243 | 6.9 | 0 | 3 | 1.1 | 1 |
| Redox cofactor | 225 | 6.4 | 0 | 2 | 1.0 | 1 |
| NAGGN | 216 | 6.1 | 0 | 1 | 1.0 | 1 |
| Arylpolyene | 177 | 5.0 | 0 | 2 | 0.8 | 1 |
| NAPAA | 142 | 4.0 | 0 | 1 | 0.6 | 1 |
| RiPP-like | 19 | 0.5 | 0 | 2 | 0.1 | 0 |
| Betalactone | 13 | 0.4 | 0 | 1 | 0.06 | 0 |
| Triceptide | 3 | 0.1 | 0 | 1 | 0.01 | 0 |
| Ranthipeptide | 1 | 0.03 | 0 | 1 | 0.00 | 0 |

BGC: biosynthetic gene cluster, NAGGN: N-acetylglutaminyglutamine amide, NAPAA: non-alpha poly-amino acids like e-Polylysine, NI: NRPS-independent, NRPS: non-ribosomal peptide synthetases, RiPP: ribosomally synthesised and post-translationally modified peptide.

To determine the similarity among the predicted BGCs and deduplicate them, we clustered all BGCs into gene cluster families (GCF) based on sequence similarity using BiG-SCAPE (38) A GCF contains BGCs that are predicted to produce chemically-related specialized metabolites. The 3,519 BGCs were clustered into 148 GCFs (Table S2). Almost one-third of the GCFs were singletons, i.e., they consisted of a single BGC found in only one genome. A GCF was present in a median of four genomes (mean = 21.6, IQR = 1-15, Figure 2C). Nine GCFs were found in over 100 isolates, predicted to produce compounds of the classes arylpolyene, hydrogen cyanide, redox cofactor, NPRS, NRPS-like, NAPAA and terpene. The two most prevalent families, which encoded a hydrogen cyanide and a redox cofactor, were almost fixed in the population, missing from only one and three genomes, respectively.

Only four GCFs had a reference specialized metabolite in the database of experimentally characterized BGCs, MIBiG (39): FAM_00334 (NRPS) had hits to biosurfactants cichofactin and syringafactin, FAM_00335 (NRPS) to virginiafactin, and FAM_00462 (NI-siderophore) to the siderophore achromobactin, all known from *Pseudomonas* species, while FAM_00458 (terpene) had hits to carotenoids from *Enterobacteriaceae* and *Pantoea* (Table S3). There was a strong and significant correlation between the number of BGCs and the number of GCFs predicted in a genome (Pearson’s correlation coefficient = 0.99, p-value < 0.0001, Figure 2D), These results indicate that there is little redundancy in the BGCs encoded by a given *P. viridiflava* isolate, as BGCs of the same class usually belong to different GCFs. In addition, most of the specialized metabolites encoded by *P. viridiflava* genomes appear to be novel and/or have not yet been experimentally characterized. Their biological functions therefore remain unknown.

### Biosynthetic genes for specialized metabolites have a fitness cost for *P. viridiflava in planta*

Microbial specialized metabolites often mediate ecologically important functions, including the interaction with their hosts and with other microorganisms. Hence, it is not unlikely that the effect of carrying a BGC on bacterial fitness varies in different contexts. In this pathosystem, bacterial load correlates positively with disease severity (27). To assess the fitness benefits or fitness costs of encoding a BGC *in planta*, we used a pool of BarSeq mutants in the *P. viridiflava* ATUE5:p25.C2 background (40), a highly virulent isolate on *A. thaliana* Ey15-2 (24).

*Pseudomonas viridiflava* ATUE5:p25.C2 putatively encodes 11 BGCs in 8 regions of the genome, with region 6 encoding two completely-overlapping (i.e., chemical hybrid) BGCs and region 7 three partially-overlapping (i.e., neighboring hybrid) BGCs (Table 2, Figure S2). These 11 BGCs contain 229 genes, with the BarSeq mutant pool having strains with mutations in 222 of these (97%). The proteins encoded by the remaining seven genes were annotated by antiSMASH 7 as hypothetical protein (region 4), transcriptional regulator LiaR, protein MbtH and glutamate tRNA ligase (region 6), and NAD-dependent glycerol dehydrogenase, transcriptional regulator LsrR and hypothetical protein (region 7).

**Table 2.** Biosynthetic gene clusters (BGCs) predicted in *P. viridiflava* ATUE5:p25.C2.

| Region | Type | Number of genes in BGC | Number of genes represented in mutant pool | DAMs Ey15-2 | DAMs Col-0 |
| --- | --- | --- | --- | --- | --- |
| Region 1 | NRPS | 32 | 32 | 0 | 6 |
| Region 2 | NAGGN | 12 | 12 | 1 | 2 |
| Region 3 | NRPS-like | 30 | 30 | 0 | 4 |
| Region 4 | Hydrogen-cyanide | 10 | 9 | 0 | 0 |
| Region 5 | Redox-cofactor | 17 | 17 | 3 | 1 |
| Region 6 | NRP-metallophore,<br>NRPS | 57 | 54 | <u>10</u> | 6 |
| Region 7 | Terpene, NRPS,<br>NI-siderophore | 60 | 57 | 9 | 5 |
| Region 8 | Hydrogen-cyanide | 11 | 11 | 0 | 1 |
| Total genes |  | 229 | 222 | 23 | 25 |

BGC: biosynthetic gene cluster, DAM: differentially-abundant BarSeq mutant, NAGGN: N-acetylglutaminyglutamine amide, NI: NRPS-independent, NRPS: non-ribosomal peptide synthetases. Underlined numbers indicate that the number of DAMs was greater than expected by chance (empirical p-value $\leq 0.05$ ).

**Figure S2.**
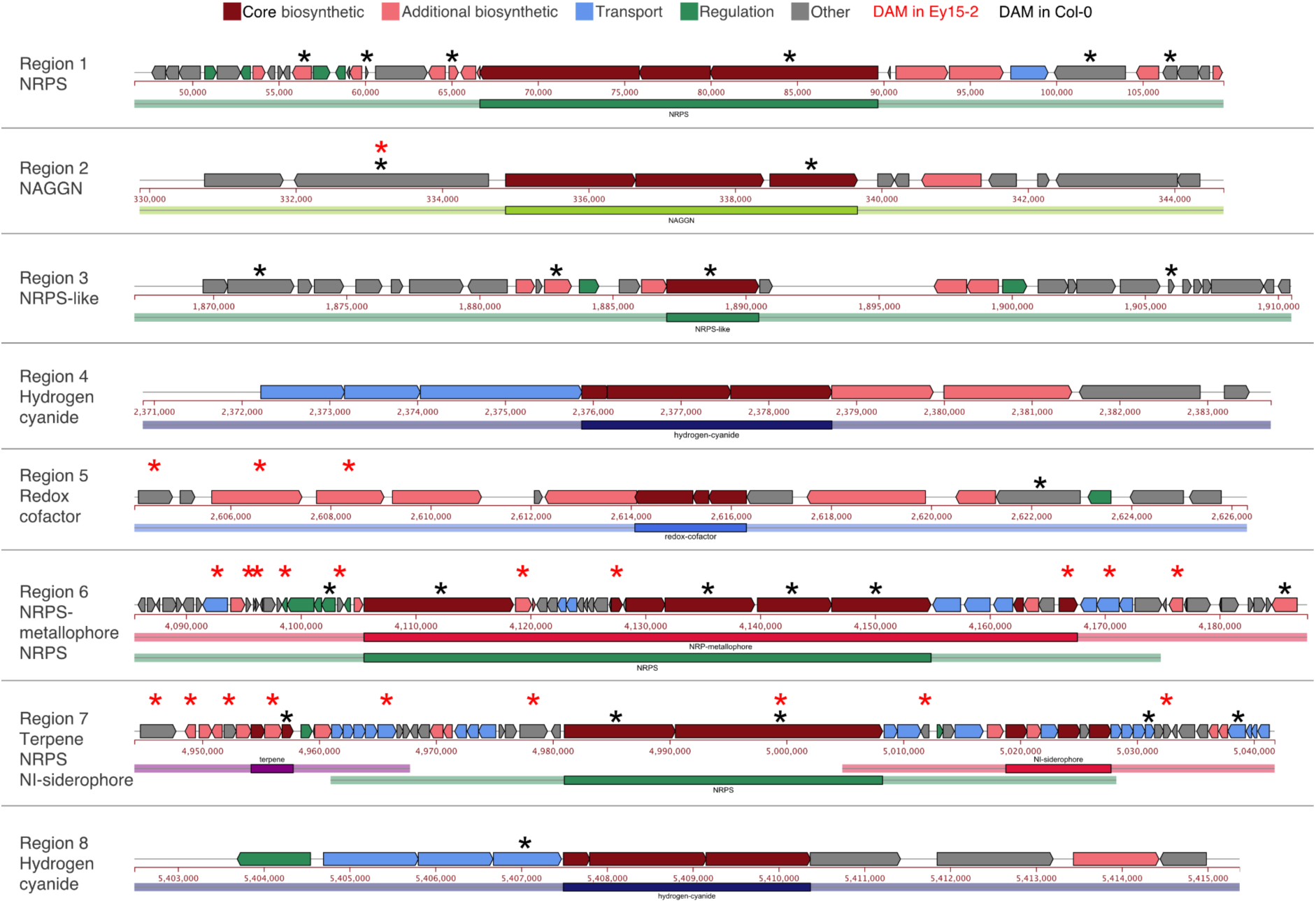
BGCs predicted in *P. viridiflava* ATUE5:p25.C2. Diagram of the eight regions encoding 11 BGCs predicted by antiSMASH. Each arrow represents a gene and the color of the arrow indicates the gene function. There were no BGCs on contigs’ edges. Asterisks on top of each BGC diagram indicate genes for which DAMs were identified in hosts Ey15-2 (red) and Col-0 (black).

We used this p25.C2 mutant pool to infect axenic *A. thaliana* Ey15-2 and Col-0 plants. RB-TnSeq (41) was applied to the pooled mutants used as inoculum and to the ones retrieved from infected plants 3 days after infection. We estimated the fitness of genes from the ratio of barcode reads in the material harvested from plants compared to the initial inoculum. We identified genes whose mutants changed in abundance in the BarSeq pool upon infection of *A. thaliana* plants with DESeq2 (42). The estimated fold change of most BGC-associated BarSeq mutants was positive across the two *A. thaliana* accessions Col-0 and Ey15-2. We identified 25 and 23 differentially-abundant mutants (DAMs) for the 222 BGC genes in Col-0 and Ey15-2 infections, respectively, defined as having a log_2_ fold change with an adjusted p-value ≤ 0.05 (Table 2, Figure 3A,B, Figure S2). To validate whether the identified DAMs represented a biological signal or a random occurrence due to the large number of genes in a BGC, we calculated the empirical p-value for each BGC region from 1,000 random sets of contiguous genes corresponding to the size of each region. The observed number of DAMs in region 6 for Ey15-2 was greater than expected by random chance (empirical p-value ≤ 0.05).

**Figure 3.**
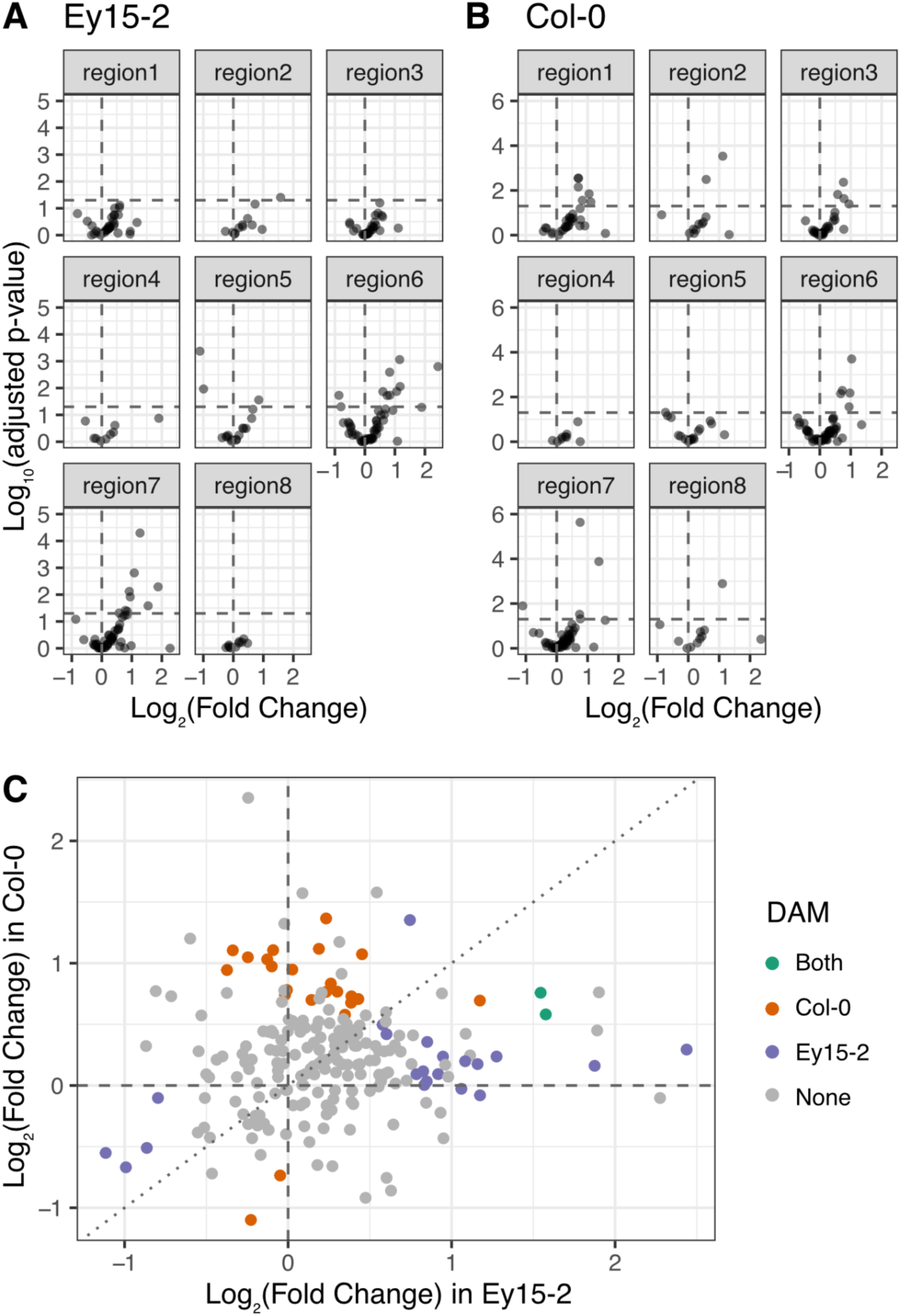
Behavior of BGC mutants *in planta*. A, B. Volcano plots of abundance change (mutant abundance *in planta* compared to inoculum) of mutants defective in 222 BGC genes, 3 days after infection of Ey15-2 (A) or Col-0 (B). Dashed lines indicate no change in abundance (x-axis, log_2_ fold change = 0) and the significance threshold (y-axis, p = 0.05). C. Correlation between abundance changes in Ey15-2 and Col-0. Colors indicate whether changes were significant in one or the other accession or in both The dotted line indicates perfect correlation. DAM: differentially abundant mutant.

**Figure S3.**
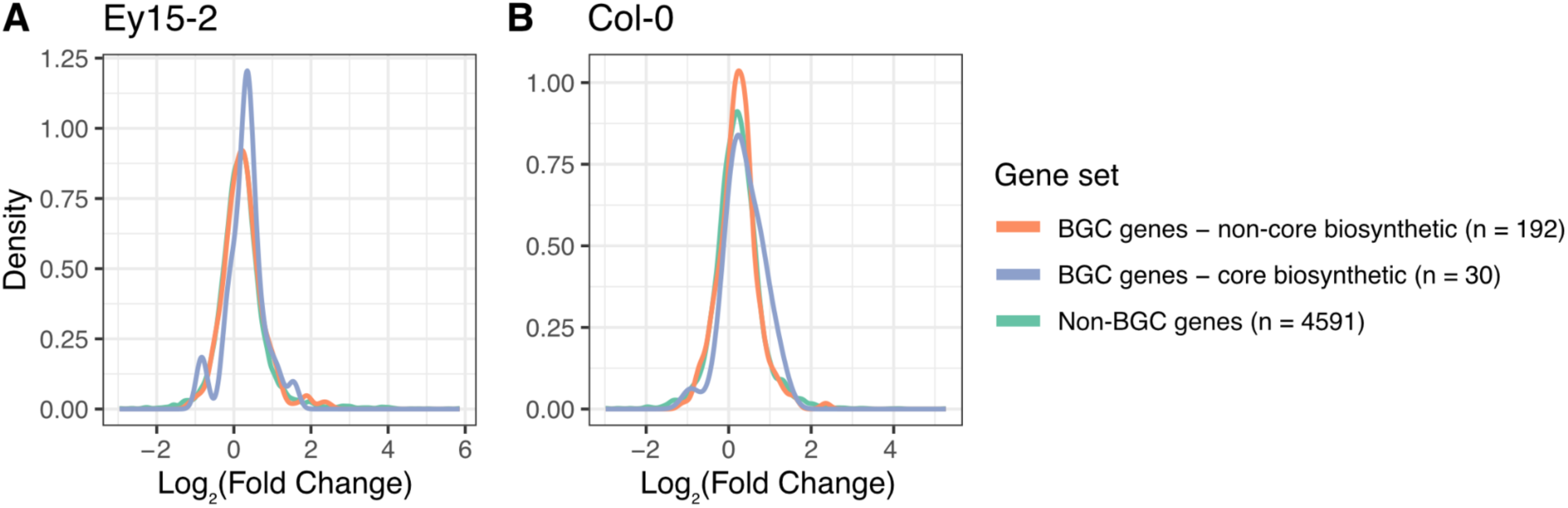
Distribution of abundance changes in the p25.C2 mutant pool. For all or a subset of mutants in hosts Ey15-2 (A) and Col-0 (B).

The longest BGCs, with the largest number of both genes and DAMs, were the 102 kb-long region 6 encoding an NRP-metallophore and an NRPS hybrid and the 97 kb-long region 7 encoding a terpene, an NRPS, and an NI-siderophore hybrid. We identified six DAMs in region 1 and four DAMs in region 3 after Col-0 infection, but none after Ey15-2 infection (Figure S2). Only two DAMs were shared between Ey15-2 and Col-0, one in region 2 and one in region 7.

We compared the fitness of each mutant in Col-0 infections against Ey15-2 infections, and found no significant correlation between the two infection environments (Pearson’s correlation coefficient = 0.07, p-value of 0.31) (Figure S3). Moreover, the direction of the fold change was not always consistent between the two accessions, with some mutants with increased abundance in Ey15-2 having a negative or no change in Col-0 and vice versa. Taken together, these results suggest that BGC fitness effects are strongly dependent on the host environment.

We were surprised to find that the abundance of most BGC mutants increased *in planta*: 74% and 68% of the BGC mutants increased in Col-0 and Ey15-2, respectively, indicating improved fitness upon BGC disruption. The abundance of only two and four mutants was reduced in Col-0 and Ey15-2, respectively. We compared the distribution of fold changes of strains with mutations in core BGC biosynthetic genes and non-core BGC biosynthetic genes (i.e., additional biosynthetic, transport-related, regulatory and other genes) as well as in non-BGC genes. The fold change of core biosynthetic genes tended to be higher than that of non-core biosynthetic genes and non-BGC genes, although the difference was not significant (Kolmogorov-Smirnov, p-value > 0.05, Figure 3D,E). Overall, contrary to our initial expectations, the increased ability of BarSeq mutants to grow *in planta* suggests that BGC production can reduce bacterial fitness.

### The presence/absence of biosynthetic gene clusters correlates with disease severity on *A. thaliana*

We had hypothesized that specialized metabolites contribute to *P. viridiflava* ATUE5 pathogenicity in *A. thaliana*. However, our RB-Tnseq experiments suggested instead that BGCs, particularly core biosynthetic genes, are more likely to carry a fitness cost rather than a fitness benefit for *P. viridiflava in planta*. Tn-seq experiments and their variants, including RB-TnSeq, measure fitness in a population-dependent context and can underestimate the fitness advantage of compounds that are public goods, i.e., products that are secreted and affect not only the producer bacterium but also everyone else in the environment. Many specialized metabolites are secreted and thus are considered public goods for members of a bacterial population that co-exist in a shared environment (43).

Because pooled mutant infections allow for cheaters to exploit secreted metabolites produced by other strains, we next tested whether BGC effects persisted when plants were infected with individual isolates. To this end, we reanalyzed an available experimental dataset where axenic *A. thaliana* Ey15-2 plants were infected with 75 *P. viridiflava* ATUE5 isolates in single-isolate infections, and plant size was measured after 7 days post-infection as a proxy for disease severity (27). In this context, only the specialized metabolites encoded by the infecting isolate are available, and thus there are no public goods from genetically different strains. The distribution of the 120 GCFs in these 75 isolates was similar to that of the entire collection, with a median of 12 GCFs per genome (mean = 13.7, min = 7, max = 30, Figure 4A). One-third of the GCFs (35%) were restricted to a single isolate. We determined the correlation between the presence of a GCF and plant size after infection, including only those GCFs present in at least 5%, but no more than 90% isolates. This resulted in 34 testable GCFs, including seven GCFs present in *P. viridiflava* ATUE5:p25.C2. For eleven GCFs, their presence correlated with plant size after infection (Pearson’s correlation coefficient between -0.546 and 0.769, adjusted p-value ≤ 0.05, Table 3, Figure 4B,C). Only one of these families (FAM_00334) had hits in the MIBiG database, to biosurfactants syringafactin and cichofactin. Pearson’s correlation coefficient was negative for two of the GCFs, suggesting that isolates encoding these specialized metabolites are more virulent on *A. thaliana* Ey15-2 than isolates that do not encode them. Three of the significant GCFs were present in *P. viridiflava* ATUE5:p25.C2. When we looked into the RB-Tnseq data for the BGC regions represented by these three families, we found these to be regions 1 (NRPS, FAM_00392), 2 (NAGGN, FAM_00470) and region 6 (NPR-metallophore-NRPS chemical hybrid, FAM_00355). Region 6 had more BGC genes that changed abundances in Ey15-2 than expected by chance (Table 2).

**Figure 4.**
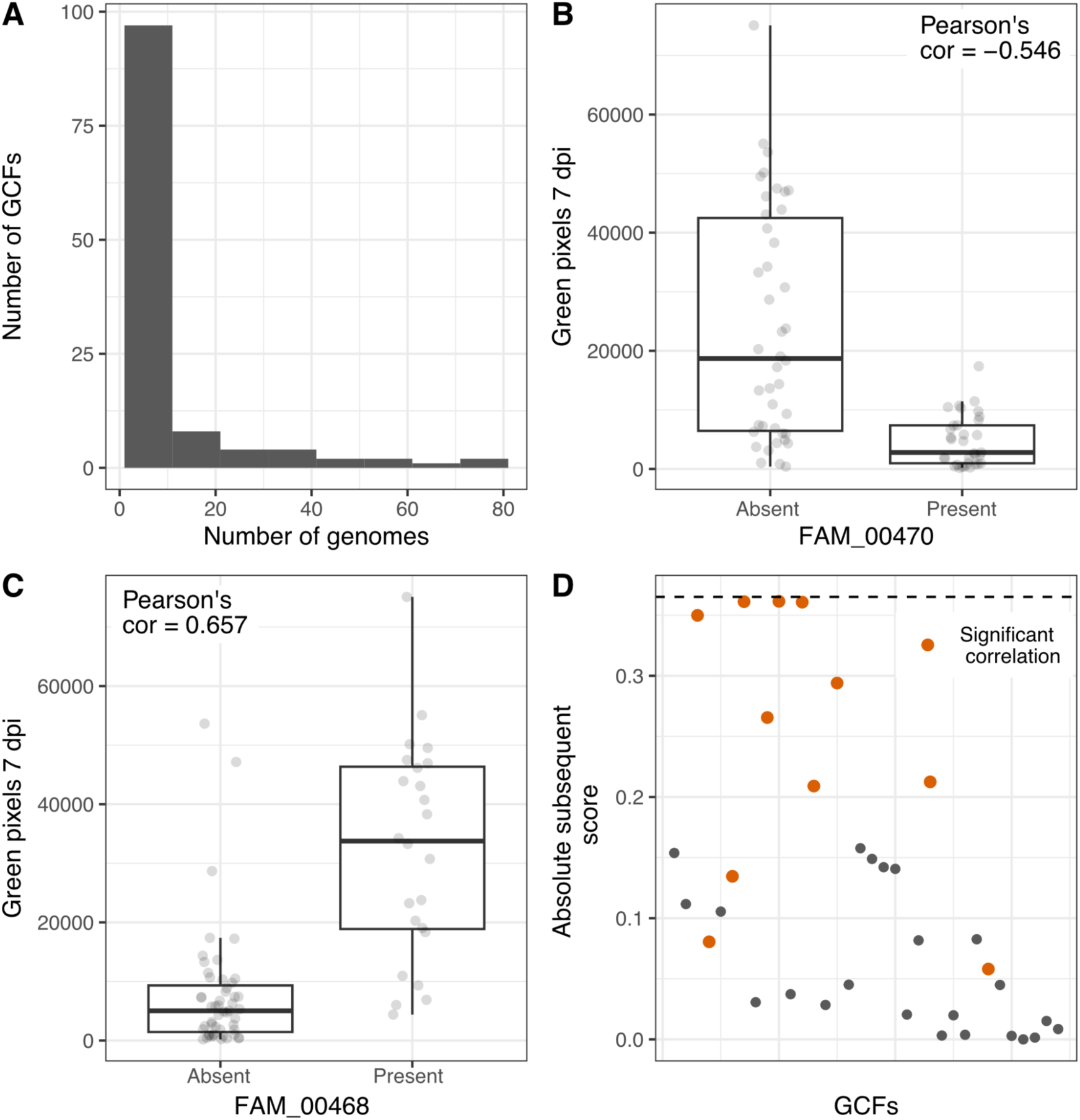
Association of gene cluster families presence/absence with disease severity *in planta.* A. Distribution of the number of genomes a given GCF is detected in a subset of 75 *P. viridiflava* genomes. B. Boxplot of the number of green pixels recorded plants 7 days after infection with *P. viridiflava* isolates where FAM_0470 was either absent or present. C. Boxplot of the plant’s green pixels 7 days after infection with *P. viridiflava* isolates where FAM_0468 was either absent or present . D. Manhattan plot of treeWAS subsequent test for 34 GCFs found in at least four but fewer than 67 *P. viridiflava* isolates. Orange points indicate GCFs whose presence/absence was significantly correlated with plant size as inferred from the number of green pixels (Table 3). GCF: Gene cluster family, dpi: days post-infection.

**Table 3.** Gene cluster families correlated with plant size after infection.

| GCF | Product class | Number of isolates (%) | Pearson's correlation coefficient | Adjusted p-value |
| --- | --- | --- | --- | --- |
| FAM_00470# | NAGGN | 33 (44) | -0.546 | 3.51x10<sup>-6</sup> |
| FAM_00392# | NRPS | 18 (24) | -0.406 | 1.45x10<sup>-3</sup> |
| FAM_00344 | NRPS | 7 (9) | 0.277 | 4.96x10<sup>-2</sup> |
| FAM_00355# | NRPS | 42 (56) | 0.278 | 4.96x10<sup>-2</sup> |
| FAM_00463 | NI-siderophore | 33 (44) | 0.312 | 2.422x10<sup>-2</sup> |
| FAM_00334* | NRPS | 20 (27) | 0.326 | 1.81x10<sup>-2</sup> |
| FAM_00384 | NAPAA | 28 (37) | 0.420 | 1.01x10<sup>-3</sup> |
| FAM_00455 | Arylpolyene | 49 (65) | 0.505 | 2.56x10<sup>-5</sup> |
| FAM_00473 | Betalactone | 11 (15) | 0.632 | 1.31x10<sup>-8</sup> |
| FAM_00468 | NAGGN | 24 (32) | 0.657 | 2.63x10<sup>-9</sup> |
| FAM_00390 | NRPS | 22 (29) | 0.769 | 2.71x10<sup>-14</sup> |

GCF: gene cluster family, BGC: biosynthetic gene cluster. \*Hits to syringafactin and cichofactin in the MIBiG database. #Families encoded by isolate p25.C2.

Population structure could influence the results of our correlation analysis, reflecting lineage rather than causality. Hence, we then applied treeWAS, a GWAS approach that corrects for population structure (44) on the same dataset of 34 GCFs in 75 ATUE5 isolates. We did not find any GCF significantly associated with disease severity with treeWAS (Figure 4D), although the GCFs with the highest absolute subsequent score were also significantly correlated with plant size. We thus cannot decisively conclude that these BGCs are virulence factors *in planta,* but they are priority candidates for functional validation.

## DISCUSSION

The genus *Pseudomonas* is known for its ability to produce many specialized metabolites (28, 29), which are often hypothesized to play a role in the interaction with other organisms and the environment. Here we show that this also applies to *Arabidopsis thaliana*-associated isolates of *P. viridiflava*, which encode a remarkably diverse and largely uncharacterized repertoire of BGCs, dominated by NRPS and NRPS-like families. Despite this extensive biosynthetic potential, RB-TnSeq experiments revealed that disrupting BGC genes generally improved bacterial growth *in planta*, indicating that specialized metabolite production often imposes a fitness cost during infection. This cost varied strongly with host genotype, suggesting substantial genotype-by-genotype interactions. Complementary analyses correlating BGC presence–absence with disease severity identified a subset of BGC families associated with altered disease severity, suggesting several candidates whose products may limit pathogenicity but be beneficial in other ecological contexts.

In our collection of 225 genomes, antiSMASH 7.1.0 (10) predicted 3,519 BGCs from 14 different chemical classes that clustered into 148 GCFs based on sequence similarity networks (38). Fragmented genomes could lead to the overestimation of BGCs due to a single BGC being distributed over multiple contigs.It is likely that the large specialized metabolite potential of *P. viridiflava* is associated with the diverse ecological niches it occupies as a generalist and opportunist pathogen, which include diverse hosts and non-plant environments (45, 46). That almost 30% of the GCFs are present in only a single isolate and that only nine GCFs are found in over 100 isolates support this hypothesis.

Fewer than 5% of the 148 gene cluster families had a closely-related hit in the MIBiG database of experimentally-characterized specialized metabolites. It is important to note, however, that even though the MIBiG database is the largest database of experimentally-validated BGCs to date, currently it contains only 3013 hand-curated BGCs and their products, 75% of which are either PKS or NRPS (39, 47). Although the use of different software and database versions makes comparisons difficult across previous studies, finding MIBiG matches for only a few BGCs encoded by the bacteria of interest seems to be the norm rather than the exception (12, 16, 48, 49). Indeed, in an analysis of over 200,000 bacterial genomes and metagenome-assembled genomes (MAGs), 97% of the predicted specialized metabolites have not yet been experimentally characterized (31).

PKS and PKS/NRPS hybrids, such as the antimicrobial mupirocin and the toxin syringolin, are known to be encoded by *Pseudomonas* isolates, including plant-associated isolates (12, 29, 30, 32, 36). However, no PKS or PKS/NRPS hybrid was predicted in the *P. viridiflava* isolates we analyzed. Half of the predicted BGCs (51%) were related to NRPS (classes NRPS, NRPS-like and NRP metallophore), in agreement with previous genome mining studies of the *Pseudomonas* genus (12, 30, 34). All isolates encoded at least three NRPS and one NRPS-like BGC, making these BGC classes the only ubiquitous ones in the genomes we analyzed.

NRPS can function as phytotoxins, antimicrobials, siderophores and surfactants, among others (50). Clustering all predicted NRPS by sequence similarity (38) resulted in 121 GCF, but only two of these families had hits to the MIBiG database (39). These hits suggest that *P. viridiflava* encodes metabolites similar to the biosurfactants syringafactin and cichofactin and to the yet-uncharacterized virginiafactin. We expect our isolates to produce cichofactin and not syringafactin, as a recent characterization of the lipopeptide diversity of the *P. syringae* species complex found that isolates belonging to phylogroups 7 and 8, which include *P. viridiflava* (51), produce cichofactin and not syringafactin (33). Although Bricout and colleagues (33) reported that isolates are either monoproducers of factins or multiproducer of the three lipopeptides families (factins, mycins and peptides), at least eight of our isolates encode BGCs closely related to both cichofactin/syringafactin and virginiafactin. The simultaneous production of both factins by a single isolate still needs to be demonstrated.

There were no hits to toxins included in the MIBiG database, confirming our previous finding that *P. viridiflava* does not encode coronatine, mangotoxin, phaseolotoxin, syringomycin, syringopeptin nor tabtoxin (24).

With antiSMASH and its predecessors, it has become increasingly straightforward to predict BGCs in microbial genomes. Tn-seq provides a complementary high-throughput approach to test the fitness of BGC genes at scale, although most of these studies have been done *in vitro*, and only a few in eukaryotic hosts (35). Between 2009, when Tn-Seq was first described, and mid-2023, fewer than 20 Tn-Seq studies with plant hosts were published (35), but 100 studies with animal hosts.

In two genetically distinct *A. thaliana* accessions, Col-0 and Ey15-2, we found that BGCs mostly reduced rather than increased *P. viridiflava* fitness. Surprisingly, the fitness costs of BGCs were strongly host dependent, and only two BGC mutants were differentially abundant in both Ey15-2 and Col-0, while the remaining 44 BGC mutants differed significantly in their abundance in only one of the two hosts. Thus, the well-documented genotype-by-genotype interactions in the *A. thaliana* - *P. viridiflava* pathosystem (27) extend also to BGCs.

We were surprised by the cost associated with BGC genes, a result that contradicted our starting hypothesis. Nonetheless, an analysis of key BGCs in *P. fluorescens*, where only 2 out of 26 Tn-seq mutants had a growth disadvantage in at least one of over 100 conditions (46, 52). As specialized metabolites can mediate not only host-microbe interactions, but also microbe-microbe and microbe-environment interactions, one potential explanation is that many of the BGCs have functions unrelated to host colonization and/or contribute to successful plant colonization by acting outside of the host, i.e., on the surrounding microbial communities. An example of this is the shift in the composition of a protective synthetic community induced by an opportunistic *Pseudomonas* root pathogen, which requires the production of two specialized metabolites, a siderophore and an antibiotic (53). This remodeling of the synthetic community contributes to an increase in root colonization by the opportunistic pathogen (53). Another possible reason for fitness cost of BGC genes is that specialized metabolites can be detected by the host and/or by host-associated microbes, triggering a defense response, and/or be toxins that kill the host and thus lead to a reduction in the bacterial counts. The *Pseudomonas* siderophore pyochelin, for example, causes a root-associated *Bacillus* isolate to increase its production of antibacterials (54).

A caveat to Tn-seq experiments is that they can lead to underestimates of fitness costs from genes that encode secreted public goods (35, 55, 56), such as siderophores, biosurfactants and toxins (57). Public goods are exploited by cheaters, individuals of the population that benefit from a public good without contributing to the production of said good, i.e., at no (fitness) cost (57). It has been shown, for instance, that mutants with defects in siderophore production have superior fitness than wild-type isolates when co-cultured, where they can cheat, but lower fitness in mono-cultures (58, 59). In addition, specialized metabolites are energetically expensive and divert resources from primary metabolism, so they can be detrimental for the producer (60) under certain conditions, similar to antibiotic resistance genes having a cost in the absence of the antibiotic (61, 62).

To overcome a possible underestimation of a BGC fitness in our RB-TnSeq experiments, we followed an orthogonal approach by examining the outcome of infections with genetically diverse *P. viridiflava* isolates that naturally vary for BGC presence. Eleven of the 34 tested GCFs correlated with disease severity in the *A. thaliana* accession Ey15-2. Similar to the results of the RB-TnSeq experiments, most GCFs were correlated with reduced disease severity, i.e., larger plant size measured in green pixels, but two were associated with increased disease severity.

We found four NRPS associated with reduced disease severity in infected *A. thaliana*, including the one predicted to be related to the biosurfactant cichofactin. This product therefore does not appear to be a virulence factor in *A. thaliana* (at least under the conditions of mono-association in an axenic system) despite previous reports (6). Meanwhile, the two GCFs negatively correlated with disease severity are potential virulence factors in *A. thaliana* infection and should be experimentally validated. One encodes a NAGGN (N-Acetylglutaminyl glutamine amide), a modified dipeptide with an important role in bacterial osmoregulation (63), and the other encodes a NRPS.

While our uncorrected correlations identified BGC associated both with increased and decreased disease severity, no significant association remained after phylogenetic correction. This could reflect limited power due to the number of isolates and/or non-random sampling. Hence, these candidate BGCs and the secondary metabolites they produce should be tested experimentally to confirm whether they play a role during *in planta* infection, particularly the two GCFs associated with increased disease severity. Although simple, a similar correlation approach identified BGCs potentially associated with *Pseudomonas* inhibition of the potato pathogen *Streptomyces scabies* (36). Further experimental characterization confirmed a role of two BGCs, a cyclic lipopeptide and of hydrogen cyanide, on the *Pseudomonas*-*Streptomyces* interaction (36).

In summary, our results challenge the common assumption that specialized metabolite production enhances pathogen success in plant hosts. Instead, while many *P. viridiflava* isolates encode well-known specialized metabolites that serve as virulence factors, such as hydrogen cyanide, the biosurfactant cichofactin and the siderophore achromobactin, we find that specialized metabolites seem to have a limited role in the virulence of *P. viridiflava* on *A. thaliana*, at least when tested in mono associations. As *P. viridiflava* is a generalist that is also found in water-related environments including rain drops, snowpack, rivers and lakes (45), there is ample opportunity for its specialized metabolites to provide fitness benefits in a non-plant context. Future experiments should include the characterization of the specialized metabolites produced by *P. viridiflava* isolates and their production -if any-upon plant infection, the validation of the fitness cost of BGCs in single-isolate *A. thaliana* infections using e.g., knock-outs as well as in co-infections, and the interaction between BGCs and host genotypes.

## MATERIALS AND METHODS

### Selection of representative *Pseudomonas* genomes

The genomes in this study were described before (24). To select representative genomes, we first assessed the completeness of each genome using BUSCO version 3.0.2 (64). Of 1,514 genomes, 1,338 had a single-copy completeness score equal to or above 95%. These were clustered using the following procedure:

1. Sort all genome assemblies by their total length, from the longest to the shortest.
2. In each iteration, set the longest assembly as the ‘representative’ of a newly formed cluster.
3. Assign members to the newly formed cluster based on the similarity of their orthology groups presence-absence profiles. Iterate over all genomes not yet assigned to a cluster, and compute the Jaccard similarity coefficient (65) between the presence-absence profile of orthology groups (obtained from PanX assignments (24)) of said genome and the cluster ‘representative’. The Jaccard similarity coefficient was calculated as:

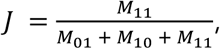

where M_11_ is the number of orthology groups shared between the strains; M_10_ and M_01_ are orthology groups present in the representative and the assessed genome, respectively. If the Jaccard similarity coefficient is greater or equal to 0.99, assign the genome to the cluster and remove it from the list.
4. Repeat steps 2 and 3 until all genomes are assigned to a cluster.

### Annotation of biosynthetic gene clusters

We used antiSMASH 7.1.0 (10) to predict biosynthetic gene clusters (BGCs) on each genome. MultiSMASH 0.3.1 was used to tabulate antiSMASH results (66). When more than one BGC was predicted in a region, each BGC was counted independently in its corresponding class. For example, a region with “NRPS, terpene, siderophore” accounted for three BGCs in our analyses. Sequence similarity networks of the BGC were generated using BiG-SCAPE (38) version 2.0.0b6, available at github.com/medema-group/BiG-SCAPE with parameters --hybrids-off and --include-singletons. We included entries from the Minimum Information about a Biosynthetic Gene (MIBiG) database (39) with parameter --mibig-version 4.0, providing manually-curated BGC annotations with known functions to the gene cluster families identified in the *P. viridiflava* genomes. The number and distribution of BGCs were plotted on the core genome maximum-likelihood tree (24) using iToL (67).

### Plant infections with pools of RB-TnSeq *P. viridiflava* ATUE5:p25.C2 mutants

BarSeq mutants were generated in *P. viridiflava* ATUE5:p25.C2 (40). Briefly, we conjugated *P. viridiflava* ATUE5:p25.C2 with the *E. coli* conjugation donor WM3064 harboring the pHLL250 mariner transposon vector library (AMD290) (68). Stocks of the mutant pool were stored at -80°C in 10% glycerol at an OD_600_ of 1.0 and were used to infect *A. thaliana*.

Col-0 and Ey15-2 wild-type *A. thaliana* plants were grown under long-day conditions (16 hours light, 8 hours dark) in an AR41L3 Percival growth chamber with 60% intensity of SciWhite LED lights. Seedlings were grown in 24-well plates on ½ Murashige & Skoog medium with vitamins and MES buffer. Thirteen-day-old seedlings were used for the infections. Plants were infected with a bacterial suspension of pooled *P. viridiflava* ATUE5:p25.C2 mutants at an OD_600_ of 0.02. Each plant was infected with 200 μL of bacterial suspension. Samples were collected in 2 mL deep-well plates three days after infection, frozen and then ground using two 5 mm glass beads. DNA was extracted using Qiagen buffers. Each biological sample contained two infected plants.

Random barcode transposon-site sequencing (RB-TnSeq) (41) was performed to determine the fitness of each mutant in the pool. Barcode sequence data from the stock mutant pool and the infected plants were obtained by multiplexing on a partial NovaSeq 6000 lane at Novogene Corporation Inc. Fitness was calculated from the abundance of reads in the plant material versus the initial inoculum pool, and analyzed with the DESeq2 package (42) in R v4.4.0 (69) to identify differentially-abundant mutants (DAMs).

To determine the statistical significance of the number of mutants for a BGC region (DAMs), we calculated the empirical p-value. For this, we generated 1,000 random sets of contiguous genes of comparable size to each BGC region and determined how many of the mutants in each random set were differentially abundant (adjusted p-value ≤ 0.05). We then calculated the probability of finding as many or more DAMs in the 1000 random sets for each region compared to the observed, i.e., experimental, number of DAMs in the same region. We used this probability as the empirical p-value.

### Correlation/association of green pixels with gene cluster families

We had previously generated data on virulence measured as green pixels of 75 genetically diverse *P. viridiflava* ATUE5 isolates on *A. thaliana* Ey15-2 (27). We calculated Pearson’s correlation coefficients between the presence/absence matrix of GCFs encoded by 5% to 90% isolates and the mean green pixels 7 days post-infection as a proxy for fitness *in planta*. The p-value was adjusted for multiple comparisons using the Benjamin-Hochberg method in R v4.4.0 (69).

Genome-wide associations between the presence/absence matrix of GCFs and green pixels after infection were obtained using treeWAS (44) in R v4.4.0(69), with 100 permutations. When a GCF had more than one copy per genome, the corresponding cell in the matrix was changed to 1 to comply with treeWAS requirements. The three association tests included in treeWAS, terminal, simultaneous and subsequent, were run. A GCF was considered significantly associated with fitness *in planta* when the adjusted p-value ≤ 0.05.

## Data availability

Genome IDs are given in Table S1. Green pixel data are from (27). Other materials and data that are reasonably requested will be made available in a timely fashion.

## ACKNOWLEDGMENTS

We thank Nadine Ziemert and Jacobo de la Cuesta-Zuluaga for their support with the specialized metabolite annotation and analysis and for their comments on the manuscript. This work was supported by the Max Planck Society, EU Horizon ERC Synergy Grant 951444 PATHOCOM (DW), USDA-NIFA 10074268 (TLK), NSF 2422727 (TLK) and NIH R35 GM150722/GM/NIGMS (TLK). Pre-submission review was conducted using ChatGPT and qed Science (https://www.qedscience.com).

ADJ, DW and TLK devised the study. HA selected the representative genomes. ADJ predicted BGCs and GCFs, ES performed RB-TnSeq experiments and their initial data analysis. MN and TLK performed single axenic infections. ADJ analyzed the data and wrote the initial draft of the manuscript. ADJ, DW and TLK reviewed and edited the manuscript.

## Notes

### Competing Interest Statement

The authors have declared no competing interest.

## REFERENCES

1. O’Brien J, Wright GD. 2011. An ecological perspective of microbial secondary metabolism. Curr Opin Biotechnol 22:552–558.

2. Etalo DW, Jeon J-S, Raaijmakers JM. 2018. Modulation of plant chemistry by beneficial root microbiota. Nat Prod Rep 35:398–409.

3. Raaijmakers JM, De Bruijn I, Nybroe O, Ongena M. 2010. Natural functions of lipopeptides from Bacillus and Pseudomonas: more than surfactants and antibiotics. FEMS Microbiol Rev 34:1037–1062.

4. Taguchi F, Suzuki T, Inagaki Y, Toyoda K, Shiraishi T, Ichinose Y. 2010. The siderophore pyoverdine of Pseudomonas syringae pv. tabaci 6605 is an intrinsic virulence factor in host tobacco infection. J Bacteriol 192:117–126.

5. Berti AD, Greve NJ, Christensen QH, Thomas MG. 2007. Identification of a biosynthetic gene cluster and the six associated lipopeptides involved in swarming motility of Pseudomonas syringae pv. tomato DC3000. J Bacteriol 189:6312–6323.

6. Pauwelyn E, Huang C-J, Ongena M, Leclère V, Jacques P, Bleyaert P, Budzikiewicz H, Schäfer M, Höfte M. 2013. New linear lipopeptides produced by Pseudomonas cichorii SF1-54 are involved in virulence, swarming motility, and biofilm formation. Mol Plant Microbe Interact 26:585–598.

7. Medema MH, Kottmann R, Yilmaz P, Cummings M, Biggins JB, Blin K, de Bruijn I, Chooi YH, Claesen J, Coates RC, Cruz-Morales P, Duddela S, Düsterhus S, Edwards DJ, Fewer DP, Garg N, Geiger C, Gomez-Escribano JP, Greule A, Hadjithomas M, Haines AS, Helfrich EJN, Hillwig ML, Ishida K, Jones AC, Jones CS, Jungmann K, Kegler C, Kim HU, Kötter P, Krug D, Masschelein J, Melnik AV, Mantovani SM, Monroe EA, Moore M, Moss N, Nützmann H-W, Pan G, Pati A, Petras D, Reen FJ, Rosconi F, Rui Z, Tian Z, Tobias NJ, Tsunematsu Y, Wiemann P, Wyckoff E, Yan X, Yim G, Yu F, Xie Y, Aigle B, Apel AK, Balibar CJ, Balskus EP, Barona-Gómez F, Bechthold A, Bode HB, Borriss R, Brady SF, Brakhage AA, Caffrey P, Cheng Y-Q, Clardy J, Cox RJ, De Mot R, Donadio S, Donia MS, van der Donk WA, Dorrestein PC, Doyle S, Driessen AJM, Ehling-Schulz M, Entian K-D, Fischbach MA, Gerwick L, Gerwick WH, Gross H, Gust B, Hertweck C, Höfte M, Jensen SE, Ju J, Katz L, Kaysser L, Klassen JL, Keller NP, Kormanec J, Kuipers OP, Kuzuyama T, Kyrpides NC, Kwon H-J, Lautru S, Lavigne R, Lee CY, Linquan B, Liu X, Liu W, Luzhetskyy A, Mahmud T, Mast Y, Méndez C, Metsä-Ketelä M, Micklefield J, Mitchell DA, Moore BS, Moreira LM, Müller R, Neilan BA, Nett M, Nielsen J, O’Gara F, Oikawa H, Osbourn A, Osburne MS, Ostash B, Payne SM, Pernodet J-L, Petricek M, Piel J, Ploux O, Raaijmakers JM, Salas JA, Schmitt EK, Scott B, Seipke RF, Shen B, Sherman DH, Sivonen K, Smanski MJ, Sosio M, Stegmann E, Süssmuth RD, Tahlan K, Thomas CM, Tang Y, Truman AW, Viaud M, Walton JD, Walsh CT, Weber T, van Wezel GP, Wilkinson B, Willey JM, Wohlleben W, Wright GD, Ziemert N, Zhang C, Zotchev SB, Breitling R, Takano E, Glöckner FO. 2015. Minimum Information about a Biosynthetic Gene cluster. Nat Chem Biol 11:625–631.

8. Dinglasan JLN, Otani H, Doering DT, Udwary D, Mouncey NJ. 2025. Microbial secondary metabolites: advancements to accelerate discovery towards application. Nat Rev Microbiol 23:338–354.

9. Medema MH, Blin K, Cimermancic P, de Jager V, Zakrzewski P, Fischbach MA, Weber T, Takano E, Breitling R. 2011. antiSMASH: rapid identification, annotation and analysis of secondary metabolite biosynthesis gene clusters in bacterial and fungal genome sequences. Nucleic Acids Res 39:W339–46.

10. Blin K, Shaw S, Augustijn HE, Reitz ZL, Biermann F, Alanjary M, Fetter A, Terlouw BR, Metcalf WW, Helfrich EJN, van Wezel GP, Medema MH, Weber T. 2023. antiSMASH 7.0: new and improved predictions for detection, regulation, chemical structures and visualisation. Nucleic Acids Res 51:W46–W50.

11. Steinke K, Mohite OS, Weber T, Kovács ÁT. 2021. Phylogenetic distribution of secondary metabolites in the Bacillus subtilis species complex. mSystems 6.

12. Saati-Santamaría Z, Selem-Mojica N, Peral-Aranega E, Rivas R, García-Fraile P. 2022. Unveiling the genomic potential of Pseudomonas type strains for discovering new natural products. Microb Genom 8.

13. Donia MS, Cimermancic P, Schulze CJ, Wieland Brown LC, Martin J, Mitreva M, Clardy J, Linington RG, Fischbach MA. 2014. A systematic analysis of biosynthetic gene clusters in the human microbiome reveals a common family of antibiotics. Cell 158:1402–1414.

14. Paoli L, Ruscheweyh H-J, Forneris CC, Hubrich F, Kautsar S, Bhushan A, Lotti A, Clayssen Q, Salazar G, Milanese A, Carlström CI, Papadopoulou C, Gehrig D, Karasikov M, Mustafa H, Larralde M, Carroll LM, Sánchez P, Zayed AA, Cronin DR, Acinas SG, Bork P, Bowler C, Delmont TO, Gasol JM, Gossert AD, Kahles A, Sullivan MB, Wincker P, Zeller G, Robinson SL, Piel J, Sunagawa S. 2022. Biosynthetic potential of the global ocean microbiome. Nature 607:111–118.

15. Sharrar AM, Crits-Christoph A, Méheust R, Diamond S, Starr EP, Banfield JF. 2020. Bacterial Secondary Metabolite Biosynthetic Potential in Soil Varies with Phylum, Depth, and Vegetation Type. MBio 11.

16. Zhang Z, Zhang L, Zhang L, Chu H, Zhou J, Ju F. 2024. Diversity and distribution of biosynthetic gene clusters in agricultural soil microbiomes. mSystems 9:e0126323.

17. Nayfach S, Roux S, Seshadri R, Udwary D, Varghese N, Schulz F, Wu D, Paez-Espino D, Chen I-M, Huntemann M, Palaniappan K, Ladau J, Mukherjee S, Reddy TBK, Nielsen T, Kirton E, Faria JP, Edirisinghe JN, Henry CS, Jungbluth SP, Chivian D, Dehal P, Wood-Charlson EM, Arkin AP, Tringe SG, Visel A, IMG/M Data Consortium, Woyke T, Mouncey NJ, Ivanova NN, Kyrpides NC, Eloe-Fadrosh EA. 2021. A genomic catalog of Earth’s microbiomes. Nat Biotechnol 39:499–509.

18. Bağcı C, Nuhamunada M, Goyat H, Ladanyi C, Sehnal L, Blin K, Kautsar SA, Tagirdzhanov A, Gurevich A, Mantri S, von Mering C, Udwary D, Medema MH, Weber T, Ziemert N. 2025. BGC Atlas: a web resource for exploring the global chemical diversity encoded in bacterial genomes. Nucleic Acids Res 53:D618–D624.

19. Wilkie JP, Dye DW, Watson DRW. 1973. Further hosts of Pseudomonas viridiflava. New Zealand Journal of Agricultural Research 16:315–323.

20. Goss EM, Kreitman M, Bergelson J. 2005. Genetic diversity, recombination and cryptic clades in Pseudomonas viridiflava infecting natural populations of Arabidopsis thaliana. Genetics 169:21–35.

21. Lundberg DS, de Pedro Jové R, Pramoj Na Ayutthaya P, Karasov TL, Shalev O, Poersch K, Ding W, Bollmann-Giolai A, Bezrukov I, Weigel D. 2022. Contrasting patterns of microbial dominance in the *Arabidopsis thaliana* phyllosphere. Proc Natl Acad Sci U S A 119:e2211881119.

22. Jakob K, Goss EM, Araki H, Van T, Kreitman M, Bergelson J. 2002. Pseudomonas viridiflava and P. syringae--natural pathogens of Arabidopsis thaliana. Mol Plant Microbe Interact 15:1195–1203.

23. Goss EM, Bergelson J. 2007. Fitness consequences of infection of Arabidopsis thaliana with its natural bacterial pathogen Pseudomonas viridiflava. Oecologia 152:71–81.

24. Karasov TL, Almario J, Friedemann C, Ding W, Giolai M, Heavens D, Kersten S, Lundberg DS, Neumann M, Regalado J, Neher RA, Kemen E, Weigel D. 2018. Arabidopsis thaliana and Pseudomonas Pathogens Exhibit Stable Associations over Evolutionary Timescales. Cell Host Microbe 24:168–179.e4.

25. Karasov TL, Neumann M, Leventhal L, Symeonidi E, Shirsekar G, Hawks A, Monroe G, Pathodopsis Team, Exposito-Alonso M, Bergelson J, Weigel D, Schwab R. 2024. Continental-scale associations of Arabidopsis thaliana phyllosphere members with host genotype and drought. Nat Microbiol 9:2748–2758.

26. Shalev O, Karasov TL, Lundberg DS, Ashkenazy H, Pramoj Na Ayutthaya P, Weigel D. 2022. Commensal Pseudomonas strains facilitate protective response against pathogens in the host plant. Nat Ecol Evol 6:383–396.

27. Duque-Jaramillo A, Ulmer N, Alseekh S, Bezrukov I, Fernie AR, Skirycz A, Karasov TL, Weigel D. 2023. The genetic and physiological basis of Arabidopsis thaliana tolerance to Pseudomonas viridiflava. New Phytol 240:1961–1975.

28. Shahid I, Malik KA, Mehnaz S. 2018. A decade of understanding secondary metabolism in Pseudomonas spp. for sustainable agriculture and pharmaceutical applications. Environmental Sustainability 1:3–17.

29. Gross H, Loper JE. 2009. Genomics of secondary metabolite production by Pseudomonas spp. Nat Prod Rep 26:1408–1446.

30. Alam K, Islam MM, Li C, Sultana S, Zhong L, Shen Q, Yu G, Hao J, Zhang Y, Li R, Li A. 2021. Genome Mining of Pseudomonas Species: Diversity and Evolution of Metabolic and Biosynthetic Potential. Molecules 26.

31. Gavriilidou A, Kautsar SA, Zaburannyi N, Krug D, Müller R, Medema MH, Ziemert N. 2022. Compendium of specialized metabolite biosynthetic diversity encoded in bacterial genomes. Nat Microbiol 7:726–735.

32. Rieusset L, Rey M, Muller D, Vacheron J, Gerin F, Dubost A, Comte G, Prigent-Combaret C. 2020. Secondary metabolites from plant-associated Pseudomonas are overproduced in biofilm. Microb Biotechnol 13:1562–1580.

33. Bricout A, Morris CE, Chandeysson C, Duban M, Boistel C, Chataigné G, Lecouturier D, Jacques P, Leclère V, Rochex A. 2022. The diversity of lipopeptides in the Pseudomonas syringae complex parallels phylogeny and sheds light on structural diversification during evolutionary history. Microbiol Spectr 10:e0145622.

34. Stringlis IA, Zhang H, Pieterse CMJ, Bolton MD, de Jonge R. 2018. Microbial small molecules - weapons of plant subversion. Nat Prod Rep 35:410–433.

35. Torres M, Paszti S, Eberl L. 2024. Shedding light on bacteria-host interactions with the aid of TnSeq approaches. MBio 15:e0039024.

36. Pacheco-Moreno A, Stefanato FL, Ford JJ, Trippel C, Uszkoreit S, Ferrafiat L, Grenga L, Dickens R, Kelly N, Kingdon AD, Ambrosetti L, Nepogodiev SA, Findlay KC, Cheema J, Trick M, Chandra G, Tomalin G, Malone JG, Truman AW. 2021. Pan-genome analysis identifies intersecting roles for Pseudomonas specialized metabolites in potato pathogen inhibition. Elife 10.

37. Anand A, Falquet L, Abou-Mansour E, L’Haridon F, Keel C, Weisskopf L. 2023. Biological hydrogen cyanide emission globally impacts the physiology of both HCN-emitting and HCN-perceiving Pseudomonas. MBio 14:e0085723.

38. Navarro-Muñoz JC, Selem-Mojica N, Mullowney MW, Kautsar SA, Tryon JH, Parkinson EI, De Los Santos ELC, Yeong M, Cruz-Morales P, Abubucker S, Roeters A, Lokhorst W, Fernandez-Guerra A, Cappelini LTD, Goering AW, Thomson RJ, Metcalf WW, Kelleher NL, Barona-Gomez F, Medema MH. 2020. A computational framework to explore large-scale biosynthetic diversity. Nat Chem Biol 16:60–68.

39. Zdouc MM, Blin K, Louwen NLL, Navarro J, Loureiro C, Bader CD, Bailey CB, Barra L, Booth TJ, Bozhüyük KAJ, Cediel-Becerra JDD, Charlop-Powers Z, Chevrette MG, Chooi YH, D’Agostino PM, de Rond T, Del Pup E, Duncan KR, Gu W, Hanif N, Helfrich EJN, Jenner M, Katsuyama Y, Korenskaia A, Krug D, Libis V, Lund GA, Mantri S, Morgan KD, Owen C, Phan C-S, Philmus B, Reitz ZL, Robinson SL, Singh KS, Teufel R, Tong Y, Tugizimana F, Ulanova D, Winter JM, Aguilar C, Akiyama DY, Al-Salihi SAA, Alanjary M, Alberti F, Aleti G, Alharthi SA, Rojo MYA, Arishi AA, Augustijn HE, Avalon NE, Avelar-Rivas JA, Axt KK, Barbieri HB, Barbosa JCJ, Barboza Segato LG, Barrett SE, Baunach M, Beemelmanns C, Beqaj D, Berger T, Bernaldo-Agüero J, Bettenbühl SM, Bielinski VA, Biermann F, Borges RM, Borriss R, Breitenbach M, Bretscher KM, Brigham MW, Buedenbender L, Bulcock BW, Cano-Prieto C, Capela J, Carrion VJ, Carter RS, Castelo-Branco R, Castro-Falcón G, Chagas FO, Charria-Girón E, Chaudhri AA, Chaudhry V, Choi H, Choi Y, Choupannejad R, Chromy J, Donahey MSC, Collemare J, Connolly JA, Creamer KE, Crüsemann M, Cruz AA, Cumsille A, Dallery J-F, Damas-Ramos LC, Damiani T, de Kruijff M, Martín BD, Sala GD, Dillen J, Doering DT, Dommaraju SR, Durusu S, Egbert S, Ellerhorst M, Faussurier B, Fetter A, Feuermann M, Fewer DP, Foldi J, Frediansyah A, Garza EA, Gavriilidou A, Gentile A, Gerke J, Gerstmans H, Gomez-Escribano JP, González-Salazar LA, Grayson NE, Greco C, Gomez JEG, Guerra S, Flores SG, Gurevich A, Gutiérrez-García K, Hart L, Haslinger K, He B, Hebra T, Hemmann JL, Hindra H, Höing L, Holland DC, Holme JE, Horch T, Hrab P, Hu J, Huynh T-H, Hwang J-Y, Iacovelli R, Iftime D, Iorio M, Jayachandran S, Jeong E, Jing J, Jung JJ, Kakumu Y, Kalkreuter E, Kang KB, Kang S, Kim W, Kim GJ, Kim H, Kim HU, Klapper M, Koetsier RA, Kollten C, Kovács ÁT, Kriukova Y, Kubach N, Kunjapur AM, Kushnareva AK, Kust A, Lamber J, Larralde M, Larsen NJ, Launay AP, Le N-T-H, Lebeer S, Lee BT, Lee K, Lev KL, Li S-M, Li Y-X, Licona-Cassani C, Lien A, Liu J, Lopez JAV, Machushynets NV, Macias MI, Mahmud T, Maleckis M, Martinez-Martinez AM, Mast Y, Maximo MF, McBride CM, McLellan RM, Bhatt KM, Melkonian C, Merrild A, Metsä-Ketelä M, Mitchell DA, Müller AV, Nguyen G-S, Nguyen HT, Niedermeyer THJ, O’Hare JH, Ossowicki A, Ostash BO, Otani H, Padva L, Paliyal S, Pan X, Panghal M, Parade DS, Park J, Parra J, Rubio MP, Pham HT, Pidot SJ, Piel J, Pourmohsenin B, Rakhmanov M, Ramesh S, Rasmussen MH, Rego A, Reher R, Rice AJ, Rigolet A, Romero-Otero A, Rosas-Becerra LR, Rosiles PY, Rutz A, Ryu B, Sahadeo L-A, Saldanha M, Salvi L, Sánchez-Carvajal E, Santos-Medellin C, Sbaraini N, Schoellhorn SM, Schumm C, Sehnal L, Selem N, Shah AD, Shishido TK, Sieber S, Silviani V, Singh G, Singh H, Sokolova N, Sonnenschein EC, Sosio M, Sowa ST, Steffen K, Stegmann E, Streiff AB, Strüder A, Surup F, Svenningsen T, Sweeney D, Szenei J, Tagirdzhanov A, Tan B, Tarnowski MJ, Terlouw BR, Rey T, Thome NU, Torres Ortega LR, Tørring T, Trindade M, Truman AW, Tvilum M, Udwary DW, Ulbricht C, Vader L, van Wezel GP, Walmsley M, Warnasinghe R, Weddeling HG, Weir ANM, Williams K, Williams SE, Witte TE, Rocca SMW, Yamada K, Yang D, Yang D, Yu J, Zhou Z, Ziemert N, Zimmer L, Zimmermann A, Zimmermann C, van der Hooft JJJ, Linington RG, Weber T, Medema MH. 2025. MIBiG 4.0: advancing biosynthetic gene cluster curation through global collaboration. Nucleic Acids Res 53:D678–D690.

40. Backman T, Latorre SM, Symeonidi E, Muszyński A, Bleak E, Eads L, Martinez-Koury PI, Som S, Hawks A, Gloss AD, Belnap DM, Manuel AM, Deutschbauer AM, Bergelson J, Azadi P, Burbano HA, Karasov TL. 2024. A phage tail-like bacteriocin suppresses competitors in metapopulations of pathogenic bacteria. Science 384:eado0713.

41. Wetmore KM, Price MN, Waters RJ, Lamson JS, He J, Hoover CA, Blow MJ, Bristow J, Butland G, Arkin AP, Deutschbauer A. 2015. Rapid quantification of mutant fitness in diverse bacteria by sequencing randomly bar-coded transposons. MBio 6.

42. Love MI, Huber W, Anders S. 2014. Moderated estimation of fold change and dispersion for RNA-seq data with DESeq2. Genome Biol 15:550.

43. Cordero OX, Wildschutte H, Kirkup B, Proehl S, Ngo L, Hussain F, Le Roux F, Mincer T, Polz MF. 2012. Ecological populations of bacteria act as socially cohesive units of antibiotic production and resistance. Science 337:1228–1231.

44. Collins C, Didelot X. 2018. A phylogenetic method to perform genome-wide association studies in microbes that accounts for population structure and recombination. PLoS Comput Biol 14:e1005958.

45. Lipps SM, Samac DA. 2022. Pseudomonas viridiflava: An internal outsider of the Pseudomonas syringae species complex. Mol Plant Pathol 23:3–15.

46. Salamzade R, Kalan LR. 2025. Context matters: assessing the impacts of genomic background and ecology on microbial biosynthetic gene cluster evolution. mSystems 10:e0153824.

47. Statistics. MIBiG Minimum Information about a Biosynthetic Gene Cluster. https://mibig.secondarymetabolites.org/stats. Retrieved 5 July 2025.

48. Mukherjee A, Tikariha H, Bandla A, Pavagadhi S, Swarup S. 2023. Global analyses of biosynthetic gene clusters in phytobiomes reveal strong phylogenetic conservation of terpenes and aryl polyenes. mSystems 8:e0038723.

49. Cimermancic P, Medema MH, Claesen J, Kurita K, Wieland Brown LC, Mavrommatis K, Pati A, Godfrey PA, Koehrsen M, Clardy J, Birren BW, Takano E, Sali A, Linington RG, Fischbach MA. 2014. Insights into secondary metabolism from a global analysis of prokaryotic biosynthetic gene clusters. Cell 158:412–421.

50. Götze S, Stallforth P. 2020. Structure, properties, and biological functions of nonribosomal lipopeptides from pseudomonads. Nat Prod Rep 37:29–54.

51. Bartoli C, Berge O, Monteil CL, Guilbaud C, Balestra GM, Varvaro L, Jones C, Dangl JL, Baltrus DA, Sands DC, Morris CE. 2014. The Pseudomonas viridiflava phylogroups in the P. syringae species complex are characterized by genetic variability and phenotypic plasticity of pathogenicity-related traits. Environ Microbiol 16:2301–2315.

52. Price MN, Wetmore KM, Waters RJ, Callaghan M, Ray J, Liu H, Kuehl JV, Melnyk RA, Lamson JS, Suh Y, Carlson HK, Esquivel Z, Sadeeshkumar H, Chakraborty R, Zane GM, Rubin BE, Wall JD, Visel A, Bristow J, Blow MJ, Arkin AP, Deutschbauer AM. 2018. Mutant phenotypes for thousands of bacterial genes of unknown function. Nature 557:503–509.

53. Amrhein A, Zhang M, Hacquard S, Heintz-Buschart A, Wippel K. 2025. Pseudomonas intra-genus competition determines the protective function of synthetic bacterial communities in Arabidopsis thaliana. PLoS Biol 23:e3002882.

54. Andrić S, Rigolet A, Argüelles Arias A, Steels S, Hoff G, Balleux G, Ongena L, Höfte M, Meyer T, Ongena M. 2023. Plant-associated Bacillus mobilizes its secondary metabolites upon perception of the siderophore pyochelin produced by a Pseudomonas competitor. ISME J 17:263–275.

55. Thibault D, Jensen PA, Wood S, Qabar C, Clark S, Shainheit MG, Isberg RR, van Opijnen T. 2019. Droplet Tn-Seq combines microfluidics with Tn-Seq for identifying complex single-cell phenotypes. Nat Commun 10:5729.

56. Schreier JE, Smith CB, Ioerger TR, Moran MA. 2023. A mutant fitness assay identifies bacterial interactions in a model ocean hot spot. Proc Natl Acad Sci U S A 120:e2217200120.

57. Smith P, Schuster M. 2019. Public goods and cheating in microbes. Curr Biol 29:R442–R447.

58. Jiricny N, Diggle SP, West SA, Evans BA, Ballantyne G, Ross-Gillespie A, Griffin AS. 2010. Fitness correlates with the extent of cheating in a bacterium. J Evol Biol 23:738–747.

59. Dumas Z, Kümmerli R. 2012. Cost of cooperation rules selection for cheats in bacterial metapopulations: Cost of cooperation and cheating in bacteria. J Evol Biol 25:473–484.

60. Drott MT, Debenport T, Higgins SA, Buckley DH, Milgroom MG. 2019. Fitness cost of aflatoxin production in Aspergillus flavus when competing with soil microbes could maintain balancing selection. MBio 10.

61. Vogwill T, MacLean RC. 2015. The genetic basis of the fitness costs of antimicrobial resistance: a meta-analysis approach. Evol Appl 8:284–295.

62. Melnyk AH, Wong A, Kassen R. 2015. The fitness costs of antibiotic resistance mutations. Evol Appl 8:273–283.

63. D’Souza-Ault MR, Smith LT, Smith GM. 1993. Roles of N-acetylglutaminylglutamine amide and glycine betaine in adaptation of Pseudomonas aeruginosa to osmotic stress. Appl Environ Microbiol 59:473–478.

64. Seppey M, Manni M, Zdobnov EM. 2019. BUSCO: Assessing Genome Assembly and Annotation Completeness. Methods Mol Biol 1962:227–245.

65. Jaccard P. 1912. The distribution of the flora in the alpine zone. New Phytol 11:37–50.

66. Reitz ZL. 2024. szreitz/multismash: v0.3.1 (v0.3.1). https://zenodo.org/records/10467162.

67. Letunic I, Bork P. 2021. Interactive Tree Of Life (iTOL) v5: an online tool for phylogenetic tree display and annotation. Nucleic Acids Res 49:W293–W296.

68. Adler BA, Kazakov AE, Zhong C, Liu H, Kutter E, Lui LM, Nielsen TN, Carion H, Deutschbauer AM, Mutalik VK, Arkin AP. 2021. The genetic basis of phage susceptibility, cross-resistance and host-range in Salmonella. Microbiology 167:001126.

69. R Core Team. 2020. R: A Language and Environment for Statistical Computing. R Foundation for Statistical Computing, Vienna, Austria.

